# Entropic Forces Drive Clustering and Spatial Localization of Influenza A M2 During Viral Budding

**DOI:** 10.1101/291120

**Authors:** Jesper J. Madsen, John M. A. Grime, Jeremy S. Rossman, Gregory A. Voth

## Abstract

The influenza A matrix 2 (M2) transmembrane protein facilitates virion release from the infected host cell. In particular, M2 plays a role in the induction of membrane curvature and/or in the scission process whereby the envelope is cut upon virion release. Here we show using coarse-grained computer simulations that various M2 assembly geometries emerge due to an entropic driving force, resulting in compact clusters or linearly extended aggregates as a direct consequence of the lateral membrane stresses. Conditions under which these protein assemblies will cause the lipid membrane to curve are explored and we predict that a critical cluster size is required for this to happen. We go on to demonstrate that under the stress conditions taking place in the cellular membrane as it undergoes large-scale membrane remodeling, the M2 protein will in principle be able to both contribute to curvature induction and sense curvature in order to line up in manifolds where local membrane line tension is high. M2 is found to exhibit linactant behavior in liquid-disordered/liquid-ordered phase-separated lipid mixtures and to be excluded from the liquid-ordered phase, in near-quantitative agreement with experimental observations. Our findings support a role for M2 in membrane remodeling during influenza viral budding both as an inducer and a sensor of membrane curvature, and they suggest a mechanism by which localization of M2 can occur as the virion assembles and releases from the host cell, independent of how the membrane curvature is produced.

**SIGNIFICANCE STATEMENT:** For influenza virus to release from the infected host cell, controlled viral budding must finalize with membrane scission of the viral envelope. Curiously, influenza carries its own protein, M2, which can sever the membrane of the constricted budding neck. Here we elucidate the physical mechanism of clustering and spatial localization of the M2 scission proteins through a combined computational and experimental approach. Our results provide fundamental insights into how M2 clustering and localization interplays with membrane curvature, membrane lateral stresses, and lipid bilayer phase behavior during viral budding in order to contribute to virion release.

## I. INTRODUCTION

Lipid membranes are macromolecular assemblies that make up the envelopes and internal compartments of living organisms. Membranes are essential because they provide physical boundaries capable of encapsulating and protecting the organisms constituents from the outside environment (1). Organization of these membranes, their constituting components and effectors, on various time and length scales is critical in many cellular processes (2) including recognition (3, 4), neuronal signaling (5–9), or endocytosis and exocytosis (10).

An infectious virus needs to cross the envelope of the host cell twice in order to complete the replication cycle – once to enter and once to exit. For this reason, any components or mechanisms related to these steps are an attractive antiviral target because they are essential for viral replication. The replication process of a virus is complex, and requires a plethora of mechanical and chemical steps. Some viruses hijack the host cell apparatus to perform certain functions; For example, the endosomal sorting complexes required for transport (ESCRT) is used by many viruses to perform tasks related to membrane remodeling. Curiously, influenza A carries its own protein called matrix 2 (M2; see Fig. 1) that is believed to facilitate the last critical step of the replication process: viral release subsequent to severing of the budding envelope (11).

**Fig. 1.**
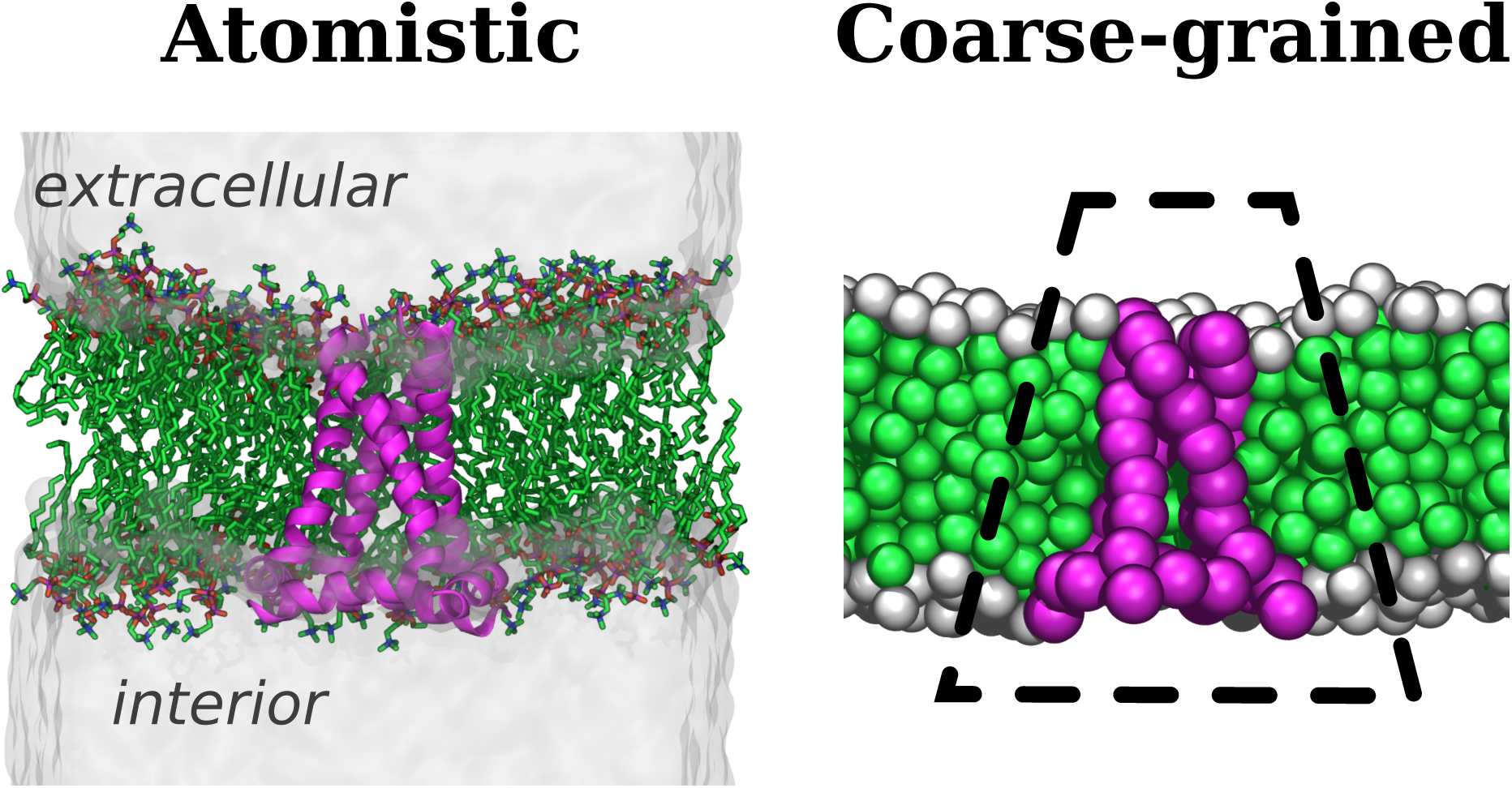
Left: The transmembrane M2 protein in an all-atom simulation of a model biological membrane. The protein is shown as purple ribbons, with membrane lipids as sticks (carbons are green, hydrogens omitted). Physiological saline is depicted as translucent blobs flanking the membrane. Right: The corresponding coarse-grained representation of the same system. The protein is depicted in purple, and the membrane lipids as white and green (for interfacial beads and tail beads respectively). M2 has a distinct conical (or “wedge”) domain topology with significant splay as indicated by the dashed line.

In this work we systematically investigate the role of M2 in budding at a coarse-grained (CG) resolution using a combination of computer simulations and experiments. The use of theory and simulation to study fundamental processes has gained wide-spread popularity in the biophysical sciences, with CG models successfully applied to the investigation of processes related to viral budding such as vesicle budding by antimicrobial peptides (12), viral proteins and capsids (13–16), dynamin constriction (17), and nano-particle endocytosis (18–21). In our simulations, theoretical as well as experimental constraints are taken into account to ensure that the system studied presents a close correspondence with biological reality. Our resultant CG model and simulations are therefore validated by comparison to experimental observations, and used to illuminate molecular details of the budding process and produce meaningful predictions.

## II. RESULTS

### The driving force for M2 cluster formation is entropic in nature and dictated by membrane lipid properties

The aggregation behavior of the M2 protein was investigated in lipid membranes of pure and mixed composition with varying lipid flexibilities ranging from soft^6ε^ to stiff^10ε^. Chemically, the soft^6ε^ lipids represent the collective behavior of highly flexible unsaturated fatty acid chain phospholipids in a liquid-disordered setting whereas stiff^10ε^ lipids represent a locally rigid composition of saturated fatty acid chain phospholipids, sterols, sphingolipids, etc. Changing the flexibility of the individual lipids will change the membrane mechanical properties in general. The dynamics of M2 (Fig. 1) in the flat tensionless membrane consisting solely of soft^6ε^ lipids are governed by normal diffusive motion. At higher M2 concentrations this naturally transitions into the sub-diffusive regime because, when embedded in the soft^6ε^ membrane, the proteins do not cluster but rather “bounce off” one another. The driving force for protein clustering under these circumstances is weaker that thermal fluctuations at room temperature (*k_B_T*) as shown in Fig. 2. However, when CG M2 is placed into a membrane consisting of the stiffer lipids, stable protein clusters spontaneously emerge. We note that such cluster formation is not irreversible, and the dynamical formation and reformation of clusters indicate that we are indeed probing true equilibrium behavior rather than merely observing kinetically trapped intermediates. The lifetime distribution of a protein-protein contacts apparently follows a power law decay with an exponent of *α =* ~2 and protein-protein contact lifetimes are therefore predicted to be scale free with no associated characteristic timescale.

**Fig. 2.**
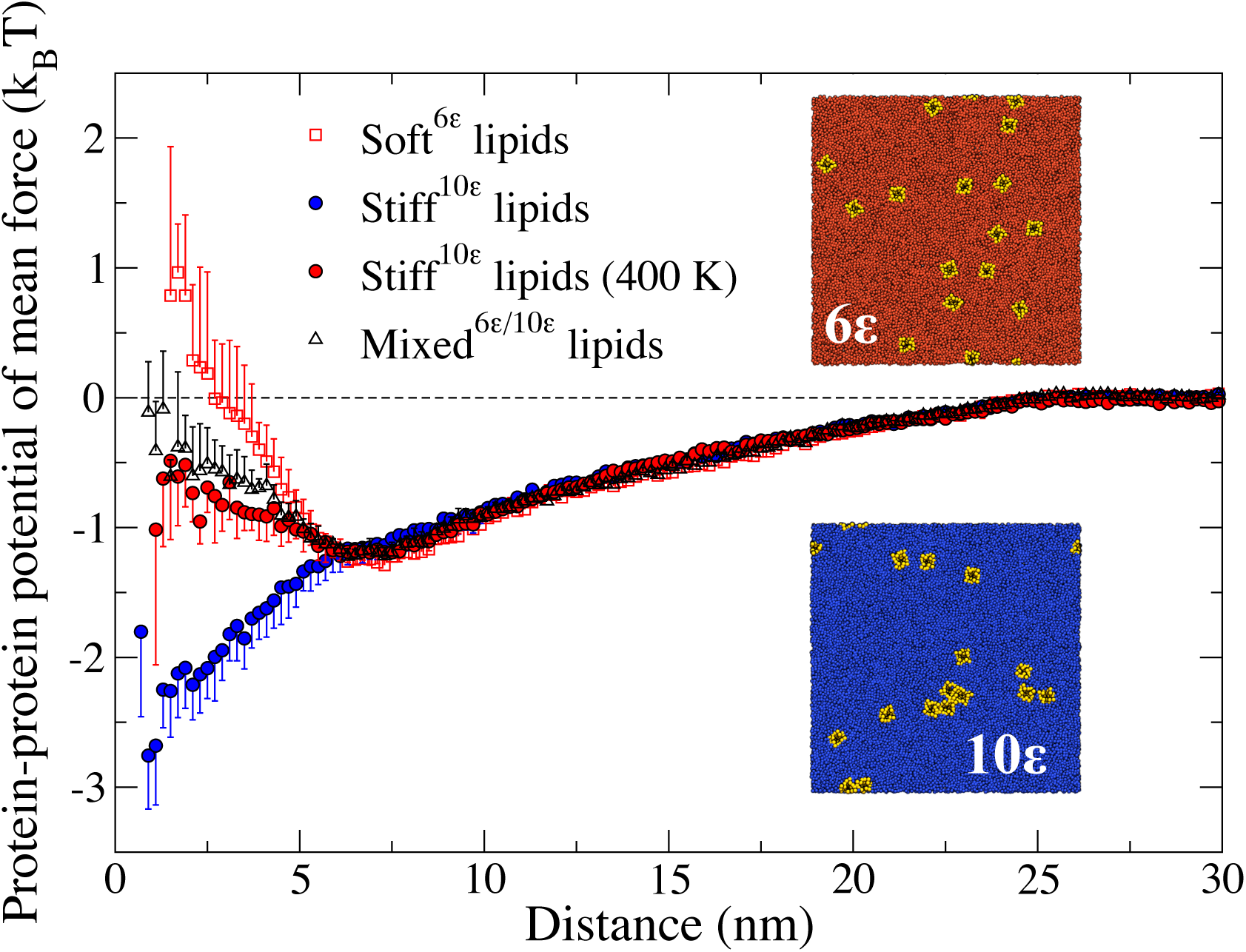
The estimated protein-protein potential of mean force for bilayer compositions of varying lipid flexibilities ranging from soft^6^*^ε^* to stiff^10^*^ε^*. M2 clustering, as accentuated in bilayers with stiff lipids, is driven by an entropic force that can be suppressed by raising the temperature of the system (compare the simulations of M2 in stiff^10^*^ε^* at 300K and 400K – blue and red filled circles, respectively). Error bars indicate the unbiased standard deviation in each 0.2 nm bin from three independent runs. Insets: Snapshots from two of the simulations showing clustering (in the system with the stiff^10^*^ε^* lipids) and no clustering (in the system with the soft^6^*^ε^* lipids), respectively.

Protein clusters will disassemble if the temperature is increased, consistent with an entropic mechanism of clustering (Fig. 2); the effective attraction between proteins (at a fixed concentration) readily reveals that any membrane-mediated clustering observed in the stiff^10ε^ membrane is suppressed upon raising the temperature of the system. As our CG model does not include any explicit protein/protein attractions beyond the renormalized hydrophobic effect inherent to the BPB model, this implies that the entropic mechanism for protein clustering emerges from the actions of the lipids in the bilayer membrane.

A system consisting of a 50/50-ratio mixture between soft^6ε^ and stiff^10ε^ lipids, mixed^6ε/10ε^, was not found to facilitate protein clustering, despite a reduction in the effective potential repulsion compared to a membrane consisting only of soft^6ε^ lipids (Fig. 2).

### M2 proteins will sort in curved membranes with and without Gaussian curvature

Clustering of the M2 protein is not localized to specific spatial regions of quasi-planar membranes. When significant membrane curvature is present, however, this situation changes because a curved bilayer membrane experiences a compression of the “inner” (most highly curved) leaflet and tension of the “outer” leaflet, around pivotal plane. Although M2 can reduce tensile and compressive stresses in the bilayer due to its conical (or, “wedge”) domain shape with significant splay, substantial protein localization is entropically disfavored. Our simulations suggest that the former effect dominates the latter, even for modest imposed curvatures, for both Gaussian and non-Gaussian curvatures (Fig. 3). The M2 distributions shown in Fig. 3 are representative for converged distributions at the entire range of physiologically relevant curvatures for the budding influenza virus (ranging from the virus diameter, *D =* ~100*nm*, to the diameter of the arrested and constricted budding neck in M2-deleted viruses, *d =* ~25*nm*); however, to facilitate convergence in the simulation, we have chosen to impose curvatures along the relevant principal axes equivalent to that of the (small) budding-neck diameter (*d* = 25*nm; κ* = 1/R = 1/12.5*nm =* 0.08*nm*^−1^). We find that for a membrane with a stable sinusoidal buckle, the M2s align with the inward-bending valley (Fig. 3, left, top view seen from the extracellular side of the membrane), which we define to be positive curvature. We observe essentially no M2 protein in flat regions of the membrane using trajectory-averaged M2 density plots (Fig. 3A, top panel). We note that proteins can, on occasion, become trapped in the negative-signed peak of the buckled membrane in an apparently metastable state.

**Fig. 3.**
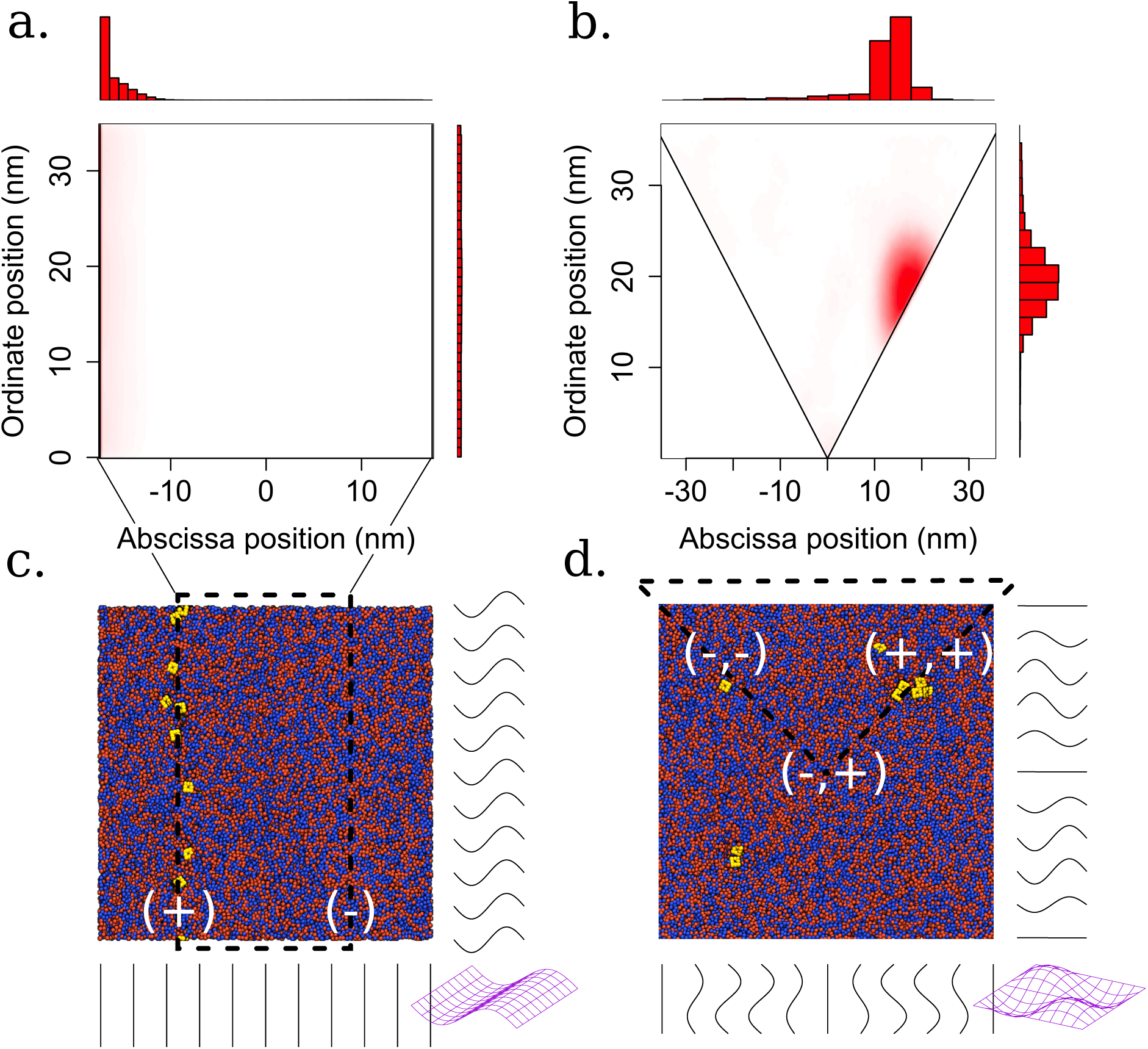
Curvature-sensing capabilities of M2 proteins in actively deformed membranes with and without Gaussian curvature. Left panels: Sampled protein density (top) in a buckled membrane. All proteins line up in the maximum (+) curvature manifold, consistent with the conical domain shape. Right panels: Protein density (top) in a membrane patch with Gaussian curvature. (−,+) = negative Gaussian curvature region, (+,+) and (−,−) = positive Gaussian curvature consistent and inconsistent with the conical M2 splay, respectively. The bordering lines in the lower panels indicate the membrane height profile along the corresponding cross-sectional cut. Imposed curvature (along the relevant principal axes) is equivalent to that of the (small) budding-neck diameter (d=25nm; x=1/R=1/12.5nm=0.08nm^−1^).

The corresponding picture in a membrane with nonzero Gaussian curvature can be seen in Fig. 3B. It is again clear curvature sorts the protein into membrane regions where its presence can reduce local stresses. M2 is found in positive curvature regions (according to our sign convention), with the most populated region where both principal curvatures are positive, denoted “(+,+)”. As seen with the buckled system, negative curvatures appear to provide a metastable protein location. Finally, we observe in the central saddle point a distinct region of negative Gaussian curvature, denoted “(−,+)”, where no significant protein density is recorded. The relative M2 populations between the region around the central saddle point and planar regions as well as the “(−,−)”-signed region are both of special interest; the former will reveal how M2 is distributed (due to curvature effects) between the neck region and the cell plasma membrane reservoir while the latter will reveal how M2 is distributed between the hemisphere of the emerging bud and the neck region. We note, however, that in the curved membrane surface with nonzero Gaussian curvature (Fig. 3B) it is hard to make a direct quantification of the mentioned relative populations from our unbiased equilibrium simulations due to the presence of the highly attractive “(+,+)” basin.

### In membranes with externally applied compressive lateral stresses, M2 clustering can induce membrane curvature to lower the free energy of the system

The envelope of a living cell is a dynamic object, responding not only to thermal fluctuations but also to mechanical and chemical influences in the surrounding environment such as scaffold proteins, osmotic gradients etc. The envelope membrane therefore experiences tensile and compressive lateral stresses under standard physiological conditions. These conditions can be mimicked in our simulations by applying external stress to the system through the barostat to investigate the effects of lateral stress on M2 clustering and membrane shape. A gentle external compressive stress of *P_ext_ =* −0.25 atm was applied to the coupled *x-* and *y*-dimensions of a simulation of 16 M2 proteins, and we observed no discernable change in behavior compared to a tensionless system for a relatively long time of ~10 M τ (Fig. 4A, region I). The size of the simulation box was essentially constant for the first 10 M integration steps (Fig. 4), but after this time M2 protein slowly localized in certain regions of the quasi-planar membrane patch (Fig. 4A, region II). After around ~20 M τ, a large protein cluster nucleates (Fig. 4A, region III and Fig. 4B, top view) with concurrent membrane curvature observed (see Fig. 4B and commensurate changes in the projected area per lipid in Fig. 4A, region III). This result is in direct correspondence with *in vitro* observations of M2 clusters in the thin necks of budding viruses (Fig. 4C), where the cell membrane is laterally stressed and surface tension is likely to be non-negligible.

**Fig. 4.**
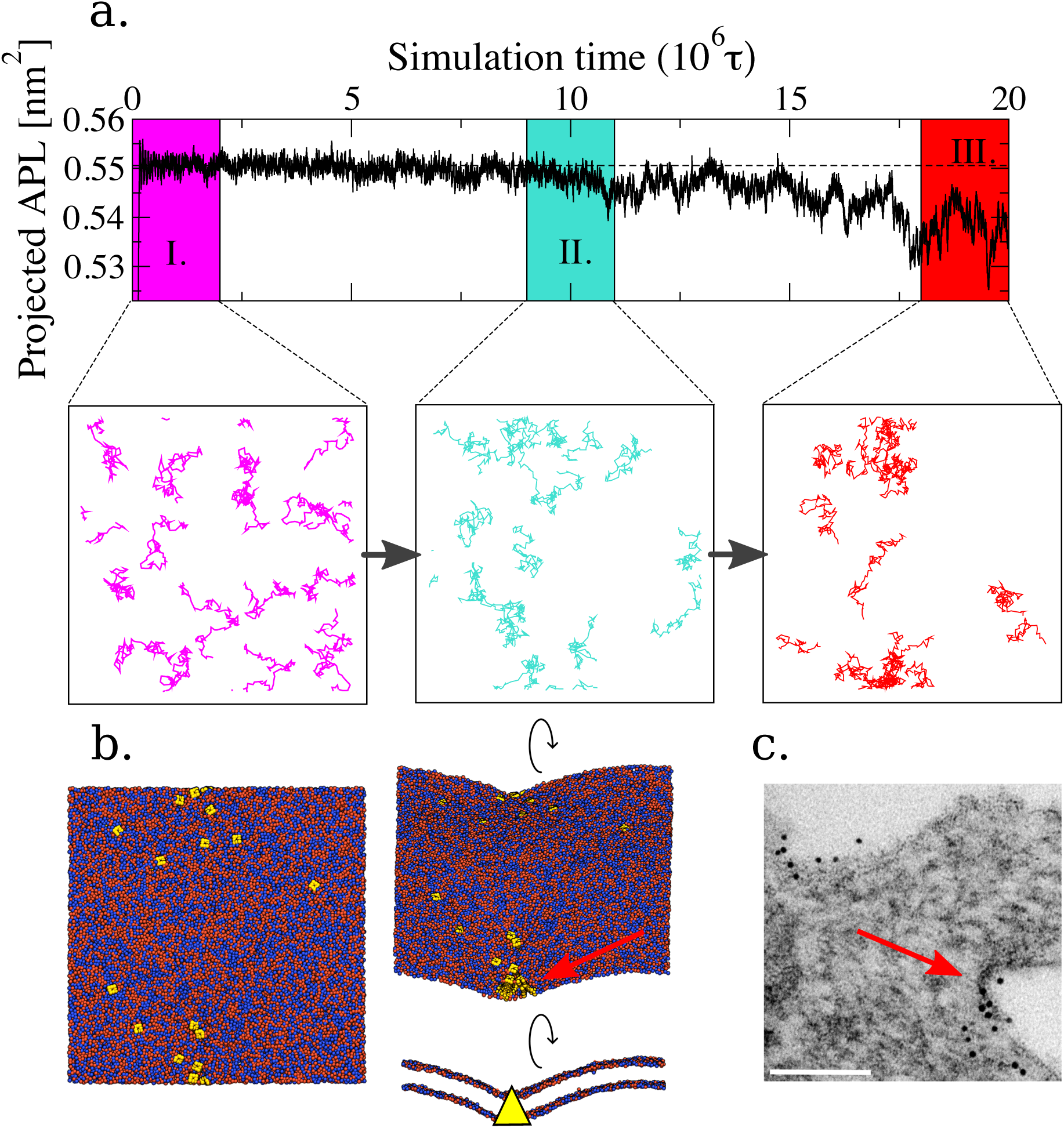
Panel A: M2 in a planar bilayer that is exposed to a constant, external compressive stress. Top: (x,y)-plane projected area per lipid in the simulation box. Three time blocks are defined for further analysis, I, II, and III. Bottom: Top view of protein tracer lines show how proteins gradually come together in the three time blocks (I, II, and III). Panel B: Final top view and side view snapshots after the membrane significantly deviates from planarity at T=20Mτ. Red arrows indicate the analogy with the in vitro system. Panel C: MDCK cells were infected with 3 pfu/cell of A/Udorn/72 for 18 hours before fixation, immuno-gold labeling of M2, thin sectioning and analysis by electron microscopy. Scale bar indicates 100 nm.

The calculated surface tension from the externally applied compression amounts to value of *γ =* −1.4 dyn/cm. However, we note that pressures calculated in CG simulations are subject to fundamental interpretation issues (22), and so this value should be considered qualitative rather than fully quantitative.

### M2 displays linactant properties in membranes with liquid-disordered/liquid-ordered (Ld/Lo) phase separation

The exclusion of M2 from the Lo phase in a phase-separated GUV provides a key insight into the mechanism of protein localization for mixed-composition membranes. Furthermore, Rossman *et al*. (11) demonstrated that M2_AH_ displays linactant properties, and this appears to be a quite general behavior for transmembrane proteins (23). We examined this effect in simulations using a mixed^6ε/10ε^ composition lipid bilayer, and introduced phase separation by weakening the cross interaction between soft^6ε^ and stiff^10ε^ lipids. This gives rise to the relative protein distributions in the Ld phase, Lo phase, and their interface as shown by the density plots in Fig. 5A. It is evident that M2 is completely excluded from the Lo phase, is freely diffusing in the Ld phase, and is slightly overrepresented at the Ld/Lo interface – exactly as indicated by fluorescently-labeled peptide corresponding to the amphipathic helix (AH) region of M2 when incubated with phase-separated GUVs (Fig. 5B) (11). Analysis of the relative fluorescent intensities along the GUV circumference shows exclusion of M2_AH_ peptide from the Lo phase, with M2_AH_ concentration at the phase boundary similarly to that observed in our CG simulations (Fig. 5A and 5B).

**Fig. 5.**
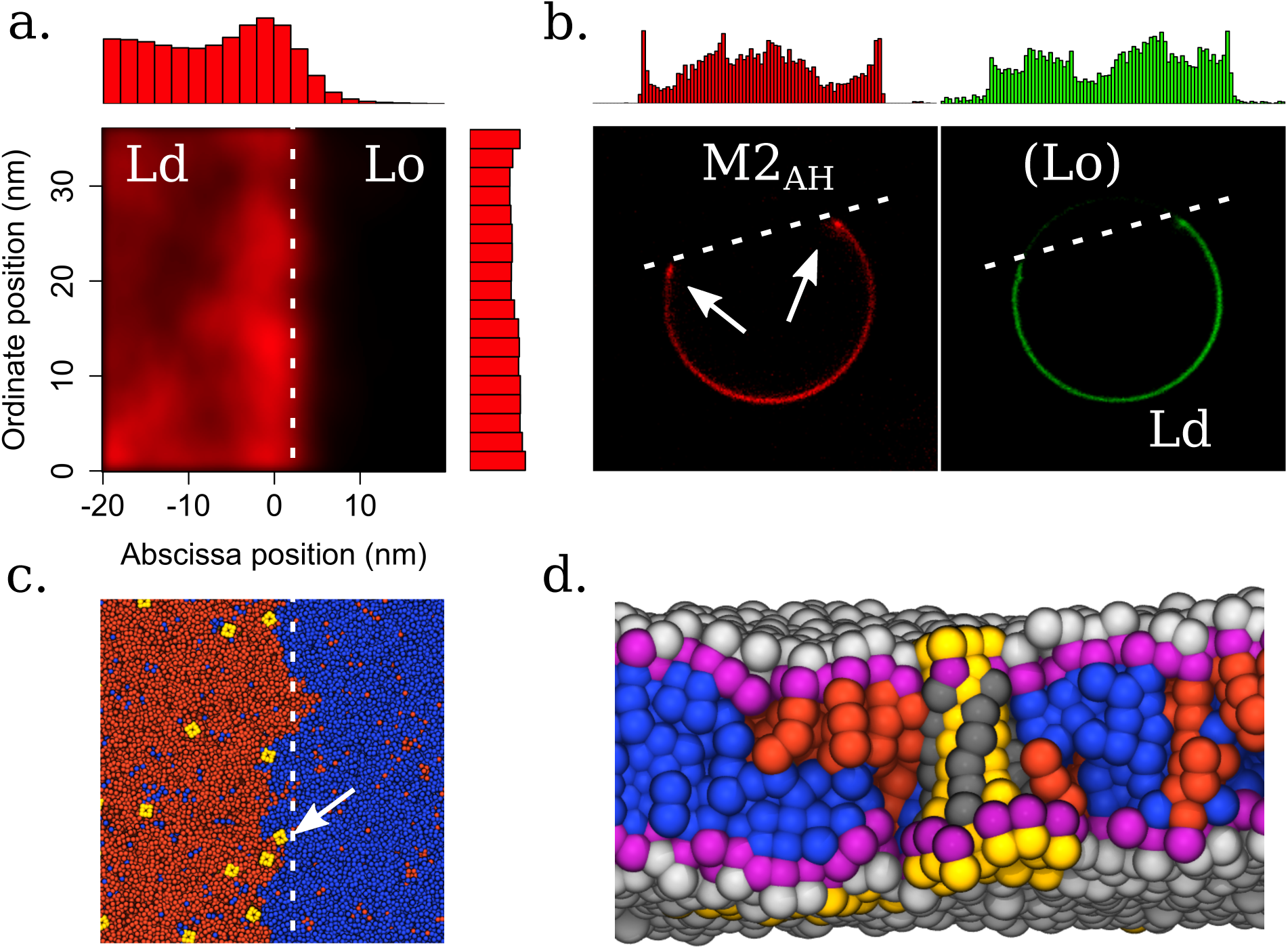
Linactant property of M2 under lipid-phase-separated conditions from simulation and experiment. Panel A: Density map of M2 positions averaged over three independent trajectories. Panel B: Representative confocal image of a phase-separated GUV, containing a fluorescent marker for the Ld phase (green) following 1 hour treatment with 10 mM of the M2_AH_-TMR peptide (red) for 1 hour. Scale bar indicates 10 μm. Images have been reanalyzed from the raw data originally presented in ref. (11). The histograms show intensity line profiles of the relative fluorescent intensity of the Ld phase marker (green) and the M2_AH_-TMR peptide (red) around the circumference of the GUV. Panel C: Simulation snapshot, top view. Blue=Lo phase, red=Ld phase, yellow=M2. Panel D: Simulation snapshot corresponding to panel C, rendered from the side in the direction of the white arrow shown in panel C. Noticeable packing defects around the protein in the Lo phase can be observed at the phase interface.

We therefore propose that the mechanism for M2’s linactant behavior is entropic depletion: M2 is excluded from the Lo phase as those lipids are too stiff to pack efficiently around the protein, resulting in the favorable exclusion of M2 protein from that phase. This effect is clearly visible in a simulation snapshot of M2 at the phase interface (Fig. 5C, arrow), where a side view (Fig. 5D) reveals packing defects of the stiff^10ε^ lipids (colored blue) in the cytosolic leaflet (bottom) around the protein. One way of analyzing the bilayer structural defects and reordering of lipids around M2 in the simulations is by comparing radial distribution functions (RDFs) of lipid and protein bead types (SI Appendix, Fig. S4). Inspection of these RDFs show that while overall number density of hydrophobic lipid beads around the hydrophobic transmembrane region of M2 appears conserved, there is a difference in the relative populations of lipids in the region corresponding to the first “solvation” shell (*r <* 10.25 Å) around the protein and this difference is accentuated by increasing lipid mixing non-ideality (i.e. upon approaching the demixing critical point) (SI Appendix, Fig. S4; e.g., compare the TAIL_M_2-TAIL_LIP/Soft_ and TAIL_M2_-TAIL_LIP_/_Stiff_ RDFs for Γ = 1.00 and Γ = 0.92).

### There exists a characteristic correlation length of soft^6ε^ lipid enrichment of M2 in mixed component bilayers that increases linearly toward the critical point of phase-separation

The same physics governing M2 protein clustering in stiff^10ε^ lipid membranes and linactant property in phase-separated soft^6ε^-stiff^10ε^ lipid membranes gives rise to a spatial correlation length wherein soft^6ε^ lipids are enriched around the M2 protein. This effect can be quantified by the fractional enrichment, Φ, of soft^6ε^ versus stiff^10ε^ lipids around each protein (Fig. 6). The fractional enrichment has exponential decay, and the correlation length, *λ*, can be retrieved by fitting to the appropriate exponential decay function. These results show that the lipids surrounding the protein clearly deviate from ideal mixing (i.e., *λ* = 0). Instead, the soft^6ε^ lipids pack preferentially around the protein even when no enthalpic discrimination exists for the lipid/lipid and lipid/protein interactions (i.e., Γ = 1.00). The trend we observe predicts that approaching the phase-separation critical point of the lipid mixture will produce an approximate linear increase in the characteristic correlation length.

**Fig. 6.**
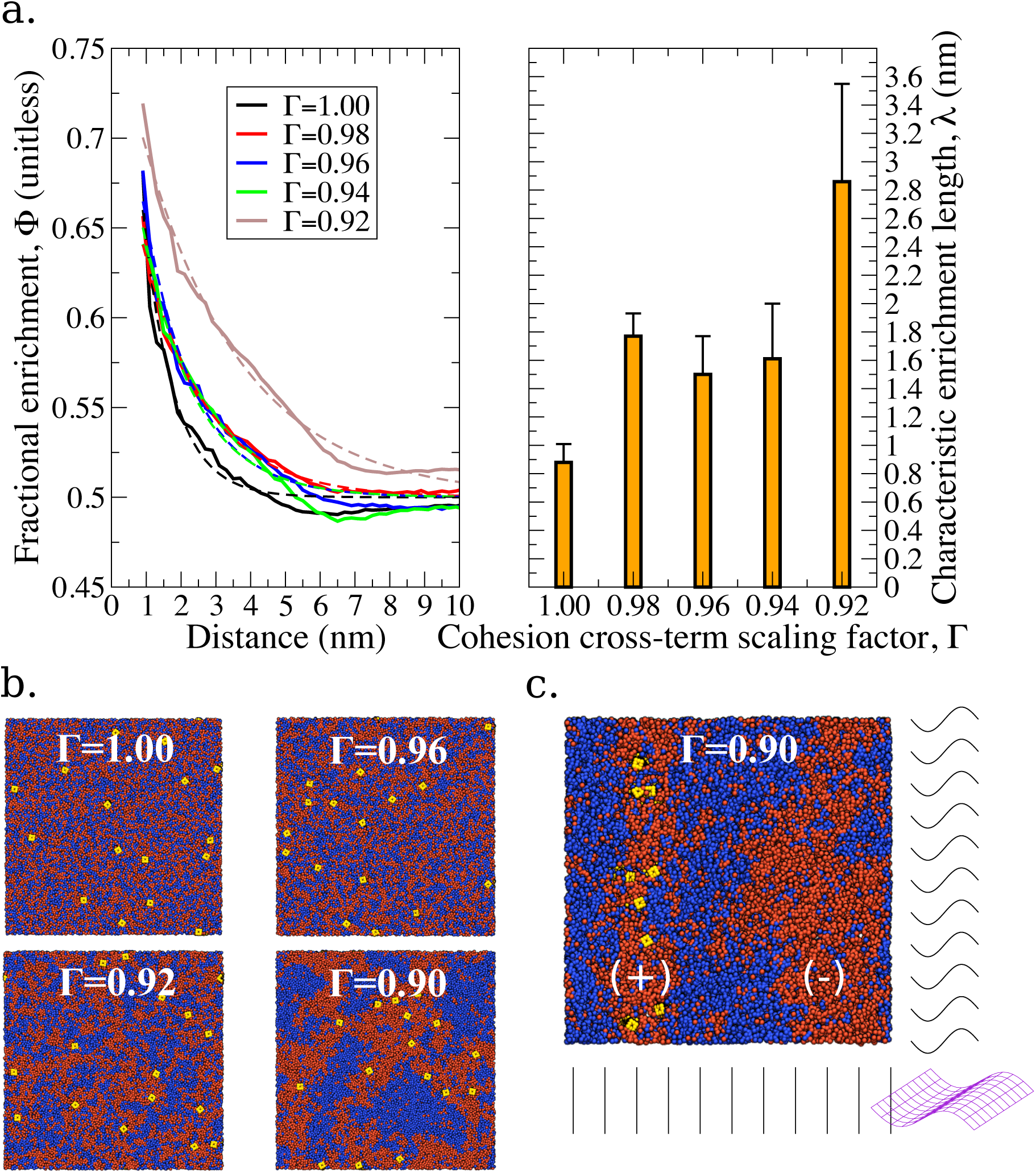
Fractional enrichment of soft^6ε^ lipids to stiff^10ε^ lipids in a mixed^6ε/10ε^ lipid bilayer membrane as it approaches its phase-separation (demixing) critical point. This is achieved by decreasing the cohesion cross-term interaction between the two lipid species of the mixture. The characteristic length, λ, associated with the soft^6ε^ lipid enrichment is estimated by a non-linear curve fitting of the fractional enrichment to a single exponential decay of the form f(x) = C_0_ * exp(-x/λ) + 0.5 (the fits are shown as dashed lines) over the entire range within 35nm. Scaling factors less than 0.92 could not be convincingly fit to this form. Error bars in the histogram show standard deviation estimated from block analysis of the trajectories. Notice how, even at Γ=1.00, there is a characteristic correlation length greater than zero showing that the lipids are non-ideally mixed around the protein. Insets: Snapshots from the simulations corresponding to Γ=[1.00, 0.96, 0.92].

### Both soft^6ε^ lipids and stiff^10ε^ lipids in mixed^6ε/10ε^ bilayer membranes are locally softened around the M2 protein, with and without effective phase separation

The incorporation of transmembrane proteins or other membrane inclusions will generally affect local lipid properties, in particular the nematic order. We find clear evidence that both soft^6ε^ and stiff^10ε^ lipids experience lower average nematic order parameters as measured from termini hydrophobic groups (Fig. 7). This lipid softening extends out to ~ 3 - 4 nm, and can therefore extend beyond the range of the fractional component enrichment in mixed bilayers (1 – 3 nm, Fig. 6). We predict that this behavior has little or no dependence on the effective driving forces of phase separation based on identical results (within calculated error) for simulations approaching the demixing critical point, Γ = [1.00, 0.98, 0.96, 0.94, 0.92].

**Fig. 7.**
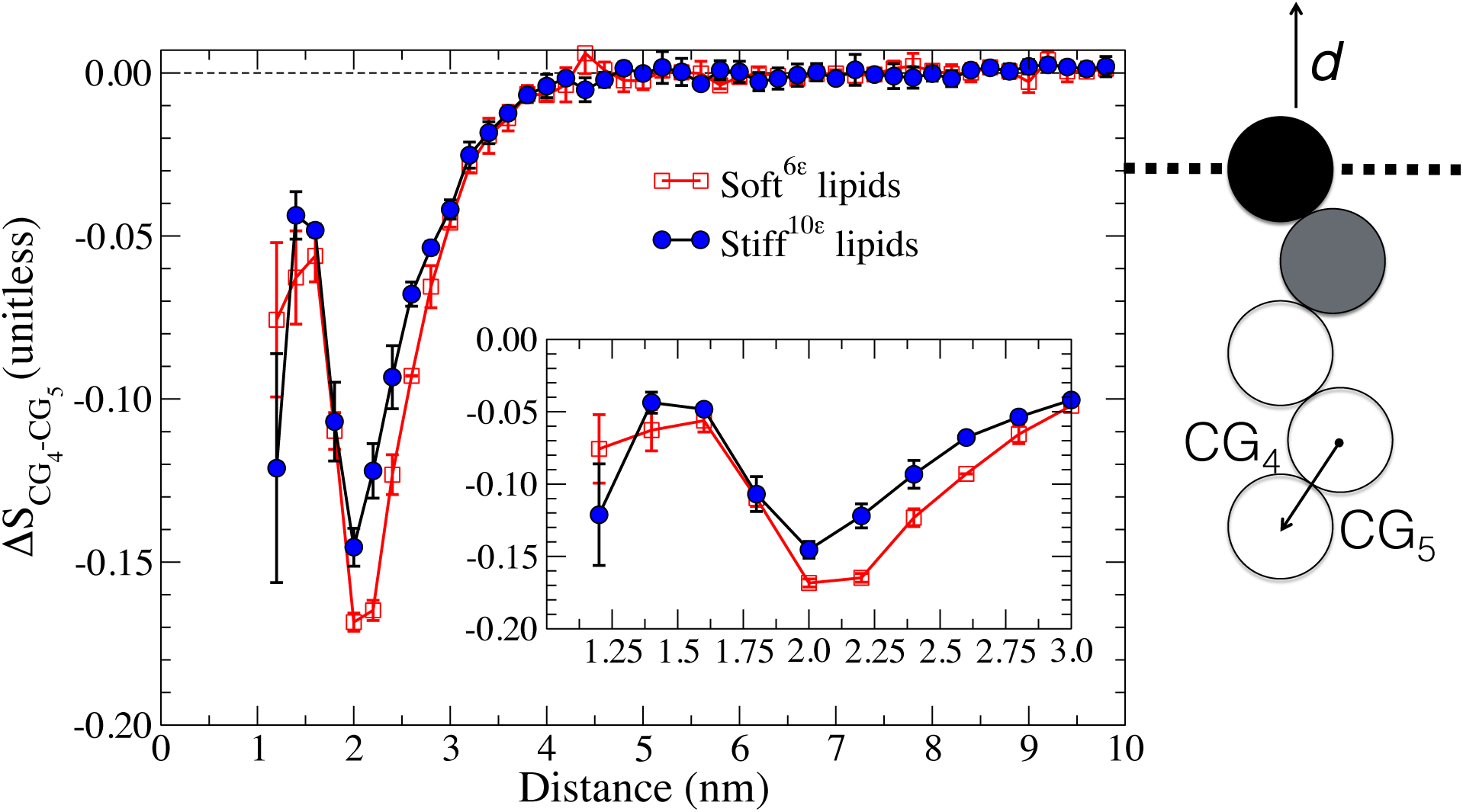
Difference in nematic order parameter of the lipid termini, ΔS_CG4-CG5_, at a given distance from M2 as compared with the value in the bulk (in infinite separation from the protein). The distance over which lipid order parameters are affected is within 4 nm and no longer-ranged effects are observed. Data is shown for the mixecf^6ε/11ε^ lipid bilatyer without the demixing term (Γ=1.00). Error bars indicate the unbiased standard deviation in each 0.2 nm bin from three independent runs. Inset: Zoomed view of the minimum well. Same units as main plot. A schematic drawing of a membrane lipid is shown on the right with the nematic director, **d**, being equal to the membrane normal.

## III. DISCUSSION

### CG model and correspondence with *in vivo* and *in vitro* systems

Our CG model of M2 is deliberately generic, and therefore its physical manifestations are in principle applicable to other transmembrane proteins under similar conditions. While recent advances have been made in the study of model influenza virions using higher resolution but still arguably ad hoc CG models (24), these models remain too computationally expensive for extensive elaborations due to the large number of degrees of freedom and long time scales required for the relaxation and adequate sampling of many important processes (milliseconds to hours). To emphasize this point consider that the state-of-the-art software and hardware used nowadays for biomolecular simulation typically is limited in practice to simulation times of tens of microseconds, and only for small system sizes. Our approach circumvents this bottleneck by integrating out (i.e., coarse-graining) degrees of freedom that can be assumed to be less relevant for the generic behavior, and we additionally benefit for accelerated dynamics in the resulting much “smoother” and lower dimensional free-energy landscape.

Necessarily, our simpler CG model comes with its own set of interpretation challenges that should now be addressed. In particular, the lipid model employed in this study should be discussed in the context of the lipids typical to cell membranes and enveloped viral particles. M2 was previously shown to play a role in the ESCRT-independent membrane envelope scission in viral release from a host cell (11), and certain molecular species such as cholesterol have been implicated in M2-dependent function (25–29). The hydrocarbon rings in cholesterol confer rigidity to the molecule (30), and a connection to cholesterol-rich micro-/nanoscale structure *in vivo* or *in vitro* can therefore be established with our stiff^10ε^ lipids on the basis of molecular flexibility in the bilayer matrix (31). While a specific interaction between M2 and cholesterol has been proposed, mediated by a putative cholesterol recognition/interaction amino acid consensus (CRAC) sequence, recent studies suggest that the actual association of cholesterol to M2 could be located in a specificity pocket between one amphipathic helix and one transmembrane domain helix (28, 32). Other evidence contradicts the idea M2 is associating with nano-domains (or so-called “rafts”) (33) and that the CRAC sequence plays a role herein (34). Nonetheless, the presence of a molecular specificity pocket on M2 toward cholesterol might provide general affinity toward the budozone area (35) or in the later stage of budding (36). While structural modulators of M2 (such as cholesterol) might affect the CG model if for instance cholesterol-binding activity can change the affinity of M2 towards the Lo budozone, it appears that M2 recruitment to the budozone is due to interactions with the matrix protein and not due to the cholesterol-binding activities. It is therefore reasonable to assume that our CG molecular-mechanical model can elucidate key behaviors of M2 by capturing the dominant physical behavior in mixed composition model bilayers where entropy is very important, and our conclusions are of direct relevance to cellular and viral membranes. With that said, our present work focuses on lipid chain flexibility and not other lipid properties (such as head group charge, average lipid shape, ability of lipids to splay and interdigitate) that might have a role in M2 clustering and localization during budding.

The M2 protein consists of a four-helix transmembrane bundle (Fig. 1). At the C-terminus of each there is an amphipathic helix (37), and recent work has shown that both M2 and the M2_AH_ amphipathic peptide can serve to facilitate budding and scission of filamentous virions (38, 39). Moreover, the M2_AH_ peptide construct retains significant function in membrane remodeling (40) and curvature sensing (41). A correspondence between models of the full-length M2 protein, the M2 construct lacking the extended C-terminal cytoplasmic tail (but including the AHs), and the M2_AH_ peptide is therefore reasonable in the study of M2’s role in budding and scission.

### Entropy-driven M2 clustering

An entropic driving force for M2 clustering based solely on lipid flexibility is elucidated in this study. We provide the minimal interaction between M2 protein and lipids required to recapitulate the known stability of M2 in lipid bilayers in order to avoid the potential excessive protein/lipid interaction strengths unduly influencing the results reported here. Calculated free-energy profiles clearly indicate a range of lipid flexibilities below which M2 will cluster (Fig. 2). We remark that this clustering is completely reversible and distinct from protein aggregation; The lifetime distribution of a protein-protein contact in a cluster follows a power law decay of exponent *α =* ~2. As a direct consequence, contact lifetimes are predicted to be scale free with no associated characteristic timescale. This effect can be rationalized by scaling arguments for a colloid depletion mechanism (42) with similarities to network edge growth (43). Furthermore, the driving mechanism M2 clustering we describe has analogues to “shape entropy” in packings of faceted particles at dense packing-ratios below the jamming transition (44).

### M2 and curvature sorting/induction

Nuclear magnetic resonance techniques have shown that M2 can induce curvature in model membranes (45). In addition, M2 can stabilize and induce negative Gaussian curvature in membrane gyroid cubic phases (Ia3d) in a wide range of protein/lipid molar ratios (46). These observations are especially intriguing because of *in vivo* observations that viruses lacking M2 will stall in the budding process prior to scission “pinch off,” giving rise to arrested structures with highly constricted budding necks of negative Gaussian curvature. In light of this, our results clarify that in fact M2 does not seem to have intrinsic preference to sort to regions of negative Gaussian curvature – instead the positive Gaussian curvature (Fig. 3B and 3D, “(+,+)” signed) appears strongly preferred.

It furthermore appears that certain principal curvatures disfavors M2 clustering; However, due to the strong tendency for M2 to sense the membrane curvature, regions outside the basins of attraction might not be sampled sufficiently to draw such conclusions. In the context of the influenza A virus budding neck, we note that curvatures present are, in our sign notation, positive Gaussian “(−,−)” curvature in the spherical part of the virion envelope, negative Gaussian “(−,+)” curvature in the budding neck and approximately flat in the cell membrane, respectively. Any observed localization of M2 the budding neck is therefore likely driven by the “wedging” mechanism by M2 along the principal axis orthogonal to the neck circumference, where M2 can “wedge” the membrane favorably, and this spatial localization of M2 is then accommodated by the formation of local membrane packing defects along the other principal axis. It is conceivable that this effect contributes to line tension in the neck and the subsequent scission process.

The induction of membrane curvature upon adding M2 in moderate copy number to a quasi-planar membrane is not observed in our model. Instead, lateral membrane compressive stresses can force M2 clustering (Fig. 4) in membrane compositions where this does not otherwise occur (e.g., in the simulations corresponding to the results shown in Fig. 2). Following M2 clustering, the membrane can buckle and deviate significantly from planarity (Fig. 4), depending on the M2 concentration and the applied membrane stresses according to the state diagram shown in the SI Appendix, Fig. S3. The specific origin of membrane stresses does not affect this finding, and we estimate that stresses on the order of those expected in physiological systems are sufficient for this process to occur. For reference, plasma membrane tension measured in cells usually ranges from 0.01–0.04 dyn/cm (47, 48), but values in the range of 0.15–0.3 dyn/cm are typical for some types of cells (49–52), which is within a factor of < 5 from the tension we apply to induce M2 aggregation and induce membrane curvature in Fig. 4 (|γ| = 1.4 dyn/cm). Recall that the external “real” and CG pressures are not in quantitative correspondence, as we remarked in a prior section, so this discrepancy might help to explain any disparity between the experimental values and those seen in the CG simulation. In simulations where tensile stresses are applied, we find that M2 cluster formation is discouraged or even suppressed, all other factors being equal. This should be interpreted as indicating the general trend of decreased *relative* cluster formation when the tensile stresses are exerted upon the membrane. In fact, this tendency could contribute the observed behavior that M2 does not appear to cluster in the plasma membrane under typical situations, where tensile stresses are present.

### Spatial organization of M2 and lipid-phase behavior in budding

Significant theoretical and experimental evidence supports the influence of lipid composition on membrane curvature, phase partitioning, and nanoscale domains in vesicular budding processes (53–66). Many of the same observations and conclusions most likely also hold in biological processes involving membrane proteins, which complicate the picture still further. The viral genome of influenza A is known to encode about 10 proteins (67), though a few additional proteins have recently been identified (68). Importantly, experimental studies of the lateral organization of these proteins during viral budding (35, 69) implicates M2, and not the usual host cell ESCRT machinery (11), in the scission of wild-type virus buds.

Our model convincingly recapitulates key experimental observations of the physical phenomena relevant for *in vitro* budding of vesicles. Most prominently, we observe the linactant property of M2 upon expulsion from the Lo phase of a phase-separated GUV and subsequent overrepresentation at the phase boundary (Fig. 5). This behavior has direct and measurable consequences for the protein-lipid interactions in lipid mixtures that (while non-ideally mixed) are not phase separated. The linactant behavior is a direct consequence of lipid properties, in particular tail flexibilities, and is sensitively distorted if for instance the soft fraction of membrane lipids is stiffened even a little (SI Appendix, Fig. S5). In Fig. 6 the fractional lipid enrichment at a given distance from M2 is quantified at various lipid phase state points approaching the phase demixing critical point. The characteristic correlation lengths we predict are well below the diffraction limit of light microscopes, but they could perhaps be measure using X-ray techniques. At around the demixing point (Fig. 6B, Γ = 0.90), larger lipids domains are starting to form, which by themselves will sort when curvature is imposed leaving preferentially soft^6ε^-lipid islands to occupy curved regions and stiff^10ε^-lipid islands to occupy flat regions of the membrane manifold (Fig. 6C). This suggests a mechanism for M2/soft^6ε^ lipid colocalization where M2 will sort to curved regions, as will soft^6ε^-lipid islands, and there is an enrichment zone of soft^6ε^ lipids around the M2 protein. In our simulations, we consistently observe that M2 localizes to curved regions much faster than any mechanism known to transport lipid islands over large spatial scale (diffusive motion of lipid islands, ripening, or nucleation and growth). This observation was not pursued quantitatively as other factors not included in our model may play an important role.

In addition, it is observed that all lipid types display lowered lipid-tail nematic order in proximity to the protein, consistent with a depletion mechanism of M2 clustering and local lipid packing defects around the protein. With the above mentioned ideas in mind it is tempting to speculate that clustering and recruitment of M2 to the budding neck of a virion in turn recruits a band of soft^6ε^ lipids, thereby increasing phase separation locally and line tension in the constricted neck to facilitate scission.

A summarizing schematic drawing depicting our proposed conceptual model of influenza budding and the role of the M2 protein can be seen in Fig. 8. Entropic sorting of flexible lipids to highly curved regions work in conjunction with soft^6ε^ lipid/M2 co-localization in order to attract M2 to the neck region from the cell plasma membrane reservoir. The neck region further attracts M2 by virtue of its positively-signed curvature principal direction and the “wedging” mechanism, while negatively-signed curvature regions are metastable or repulsive for M2. Lipid-phase behavior, in particular the linactant property of M2, discourages additional recruitment of M2 to the upper hemisphere of the emerging virus since it is enriched in stiff^10ε^ lipids at this stage.

**Fig. 8.**
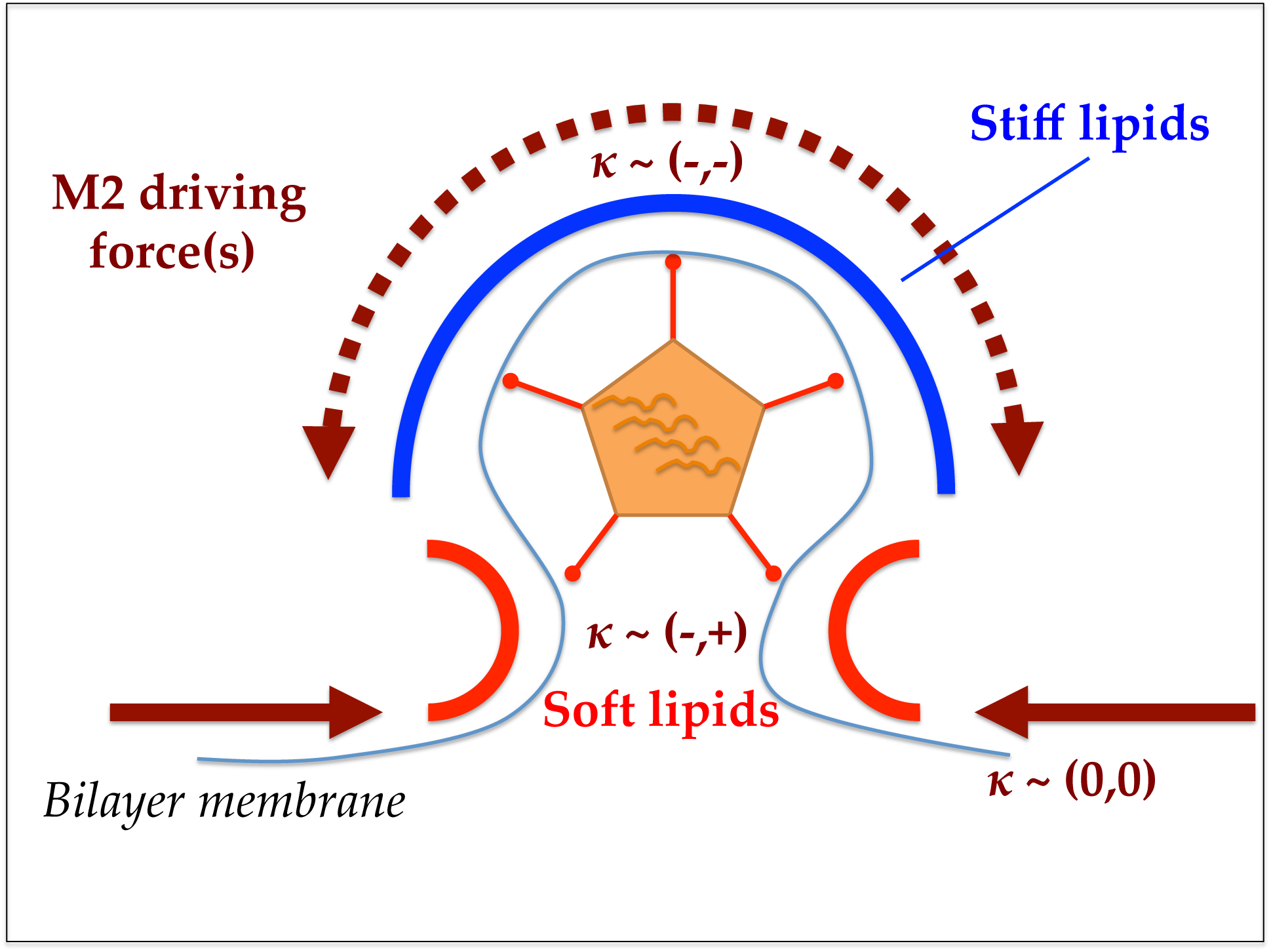
Schematic drawing of the proposed model the role of M2 in influenza budding. The entropic M2 driving forces as elucidated by our simulation results are drawn with arrows (brown). Contributing factors include partial exclusion of M2 from spatial regions where the membrane is rich in stiff lipids (due to its linactant behavior), such as the upper hemisphere (blue region) that will eventually become the mature virus envelope; and effective colocalization between M2 and soft lipids present in the neck region of the bud (red region) due to entropic and shape effects. Sign convention used for curvatures (x) are labeled according to region.

## IV. CONCLUSIONS

The present work elucidates the entropic driving forces that determine the clustering and spatial localization of M2 protein in flat and curved membranes, in membranes exposed to physiological membrane stresses, and in membranes with non-ideal lipid mixing and phase separation. These membrane conditions are prototypical to viral budding in influenza A, and our results provide new conceptual and molecular-level insights into the role of M2 in viral budding. Our coarse-grained model naturally explains key experimental observations of M2 clustering, curvature sorting in budding viruses, curvature induction in stressed membranes, and, finally, how these behaviors change depending on the collective behavior of membrane lipids. We make testable predictions regarding several relevant aspects of the physical behavior of M2, specifically: the correlation length of soft lipids and lipid softening around the protein; scale-free lifetime distribution of M2 clusters; a strong preference for positive curvatures (with negative curvature being a metastable region); and estimates of the critical M2 cluster size required to induce curvature under applied membrane stress. Our model is generic and we therefore expect our conclusions to be at least partially transferable to comparable systems in biology and soft matter physics, for example the budding of other viruses where M2-like proteins are present (e.g. influenza B) and endocytotic processes.

## V. MODELS, METHODS, AND MATERIALS

### CG Lipid Model

We have implemented a version of the tunable CG lipid model proposed by Brannigan, Phillips, and Brown (BPB) (70, 71) in the LAMMPS software (72). The BPB lipid is an implicit solvent model using 5 CG sites per molecule, and self-assembles into bilayer membranes whose mechanical properties can be tuned through the angular potential coefficient, *c_angle_* (70). The BPB model features bond and angle interactions

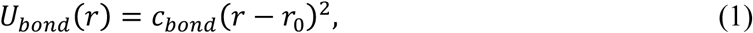

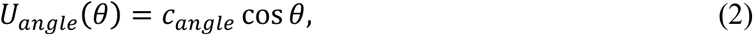

where *c_bond_* and *c_angle_* are positive constants to scale the bond and angle energies and *θ* is the (identical) angles of the lipid molecule. The model further has the following non-bonded 12-6 Lennard-Jones and long-ranged *r*^−2^ interactions

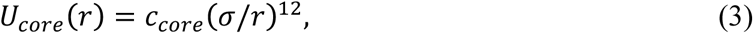

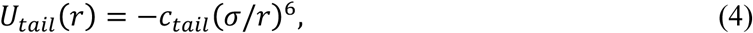

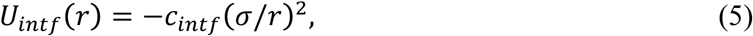

where *c_core_, c_tail_*, and *c_intf_* are positive energy constants. Interaction potentials are truncated and shifted at 2*σ* for the core and tail interactions and 3*σ* for the interfacial interaction. The values k*_B_T* =0.85*ε*, *c_core_ =*0.4*ε*, *c_tail_* = 1.0*ε*, *c_int_* = 3.0*ε* and *c_bond_* = 44*ε* and c*_bend_* = (6.0 – 10)*ε* were used as a base. We note here that our implementation differs from the original model (70) in two important aspects; lipid beads are connected with harmonic bonds rather than rigid constraints, and the effective size of the excluded-volume-only head bead type (toward the hydrophobic bead type only) was scaled by a factor of 1.25. The former was done to avoid solving rigid body constraints when running dynamics simulations, while the latter was found to discourage lipids from leaving the membrane at the target lipid flexibility given by the angle potential coefficient in the range of *c_bend_ =* (6.0 – 10)*ε*, which can be an issue (70), particularly for curved membranes in our experience. Neither modification resulted in any appreciable changes in the physical properties of the CG lipid bilayers membranes. The CG bead masses were assigned by evenly distributing the mass of the prototypical membrane lipid, 1-palmitoyl-2-oleoylphosphatidylcholine (POPC), over the 5 beads in the lipid molecule. In the relevant simulations, an energetic penalty for disfavoring cross-interactions between two lipid types can increase non-ideality and eventually cause de-mixing of the two lipid phases. This can be done either by penalizing well depth of the interaction potential that acts between the two lipids (*ε* parameter in the Lennard-Jones cross-term) or by altering the range over which the interaction is nonzero (by means of introducing a potential form with a variable range (73)). We chose to introduce a variant of the former where the interaction potential acting between tail beads of A and tail beads of B is scaled by a constant, Γ, so that the relevant potential becomes *U_tail_*(*r*) *= −*Γ*c_tail_*(*σ/r*)^6^. The modified interaction was mapped onto the tail beads, as opposed to for instance the interfacial cohesion, because this approach follows chemical intuition and was found to enable reasonable leaflet registration in curved membranes.

### All-atom MD Simulations

An atomistic structure of the M2 protein (PDB ID: 2L0J) (37) was prepared in a lipid bilayer patch as described in detail elsewhere (74, 75). Briefly, the system was constructed by the CHARMM-GUI generator (76) and simulated with molecular dynamics (MD) in NAMD2 (77) using the CHARMM36 force field (78) with CMAP (79) and the TIP3P water model (80). After an extended equilibration protocol (74) of ~100 ns duration, the system was simulated for another 100 ns to use in the parameterization of the CG model as described in the following subsection (see Fig. 1).

### CG Protein Model

A CG model of the M2 protein was constructed by mapping on average 4.1 amino acid residues into one CG site (bead), producing a highly coarse-grained model that retains the key structural features of the alpha-helical protein while maintaining a vertical resolution compatible with the CG lipid bilayer membrane (see Fig. 1 and the SI Appendix, Table S1, for the precise CG mapping). The approximate four-fold symmetry (C_4_) of the M2 homotetramer around the major axis was previously shown to enable simple construction of CG mappings (81). Non-bonded protein/lipid interactions were based on the existing BPB interactions to ensure that flanking regions of the protein (corresponding primarily to charged and polar residues) preferentially locate in the appropriate interfacial region of the bilayer (type ‘INTF’) and that the transmembrane region of the protein (corresponding to primarily hydrophobic residues) retains direct chemical compatibility with the hydrophobic beads of the bilayer interior (type ‘TAIL’). The resulting M2 model has 40 mapped CG beads in the order (INTF-TAIL_7_-INTF_2_)_4_. Purely repulsive CG beads were introduced into the central space inside the M2 channel (and also between adjacent helices) in order to maintain the overall CG protein fold and to prevent accidental lipid penetration into the hydrophilic core of the M2 channel. All membrane-flanking beads were capped with additional excluded volume beads (type ‘HEAD’) at an offset distance along *z* of *σ* to be consistent with the BPB lipid’s first bead representing the solvent above the interfacial bead in the lipid (SI Appendix, Fig. S1). The mass of the protein (5020 Da) was divided evenly among the constituting beads for simplicity and the coordinates of the model were scaled in order for the protein to span the membrane without hydrophobic mismatch. The CG bead locations were used to construct an elastic network model (ENM) to reproduce the Gaussian fluctuations recorded in the reference all-atom simulation (82, 83), with harmonic bonds introduced using a cutoff distance of 2σ (15 Å). Resting bond distances were discretized using bin-widths of 0.25 Å in order to reduce the total number of unique bond definitions required to describe the ENM, with per-residue fluctuations of the reference all-atom simulation satisfactorily reproduced using a uniform spring constant of 1.0 kcal/mol/Å^2^ for all harmonic bonds in the protein (SI Appendix, Fig. S2). The final CG M2 model has 96 beads (three bead types), and 633 bonds (38 bond types) per protein (see the supporting material for a detailed example input).

### CG Simulation Procedures

Simulations were performed with the LAMMPS MD software package (72), using the Nosé-Hoover extended Lagrangian formalism for controlling both pressure and temperature. In all cases, extremely weak coupling to thermo- and barostats were used (coupling times of 150,000τ and 15,000τ) in order to minimize perturbations of the evolution of the system. The thermodynamic ensemble sampled was constant *NPT* (constant copy number, *N*, pressure, *P*, and temperature, *T*) at *T =* 300 *K* and *P =* 0 *atm*, unless otherwise stated. We report “CG time” in units of *τ* because of the fundamental challenge in rigorously connecting simulated time with real time in CG simulations (see, e.g., ref. (84)). Simulations were run in triplicates by assigning different initial momenta distributions via distinct pseudorandom number generator seeds.

### CG System Setup

Bilayer geometries were initialized by placing CG lipids on a regular lattice in the x-,y-plane to create a membrane patch of 20,000 lipids of simulation box length ≈ 70 × 70 nm in the membrane plane. M2 proteins (if present) were embedded in the membrane with major axis aligned to the membrane normal, and overlapping lipids were removed. The number of M2 proteins inserted (up to 16, depending on the specific simulation) corresponds to an area coverage of ≈ 0% to 6% with the area per protein taken as 20 nm^2^; this somewhat exceeds the estimated average M2 coverage of an influenza virion but nonetheless provides a reasonable local range (85, 86). Membrane curvature was induced by defining repulsive shape-guiding regions (two cylinders of radius 10σ for the buckled membranes, four spheres of radius 10σ for membranes with Gaussian curvature) via harmonic exclusion potentials of the form (*fix wall/region* command)

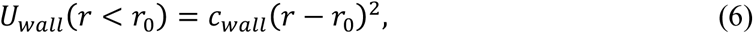

where *r* is the radial distance from the shape-guiding region. The values *c_wall_ =* 1.0 *kcal/mol* (= ~0.70*ε*) and 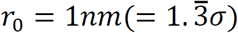 were used.

### Error Analysis

Error analysis was performed on independently executed simulation trajectories (triplets) or by an appropriate block-averaging procedure (87) of a single trajectory. Error bars show the calculated standard deviation.

### Analysis, Visualization and Plotting

Visual Molecular Dynamics (88) and in-house scripts were used for analysis and visualization, Grace (xmgrace; http://plasma-gate.weizmann.ac.il/Grace) and gnuplot (http://gnuplot.sourceforge.net) for plotting, *R* software (89) for analysis and plotting, and Inkscape (version 0.91; http://www.inkscape.org) for figure layouts.

### Lipid-Tail Nematic Order Parameter Calculation

The lipid order parameter, *S*, is given by the expectation value of the second Legendre polynomial in cos *θ* (90, 91),

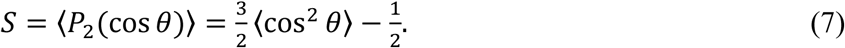

### Surface Tension Calculation

We calculate the surface tension, *γ*, by taking advantage of the three diagonal elements of the stress tensor (*σ_xx_, σ_yy_, σ_zz_*) (92),

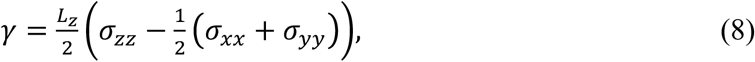

where the components of the stress tensor are generated by LAMMPS (72) and *L_z_* is the simulation box length in *z* dimension normal of the membrane plane.

### Potential of Mean Force Calculation

The Helmholtz (Gibbs) free energy was determined by the reversible work theorem according to the equation

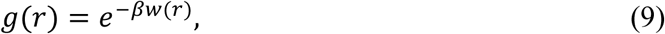

where, in the constant NVT (NPT) ensemble, the free energy for a reversible process is equal to the potential of mean force, Δ*F = w*(*r*).

### Experimental Methods

#### Giant Unilamellar Vesicles (GUVs)

Phase separated GUVs were electroformed at 60°C using a 4:4:1 molar ratio of DOPC:SM:Cholesterol (Avanti Polar Lipids, Alabaster, AL), incorporating 0.5 mol % of Bodipy-PC (Avanti) before treatment with tetra-methyl-rhodamine labeled M2 amphipathic helix (M2_AH_) peptide (GenScript, Piscataway, NJ) and confocal imaging on the LSM5 Pascal (Zeiss, Thornwood, NY) confocal microscope, as previously described (11). Image adjustment was limited to cropping and equal adjustment of levels. Line intensity histograms of Ld and M2_AH_-TMR intensities along the circumference of the GUV were calculated using the histogram feature of the Zen microscopy program (Zeiss).

#### Electron Microscopy

MDCK cells were infected as indicated and subjected to pre-embedding immunogold labeling of the M2 protein using the 14C2 mAb (85) followed by thin section processing as previously described (93). Sections were imaged on a JEOL 1230 (Tokyo, Japan) electron microscope.

## ACKNOWLEDGEMENTS

This research funded in part and the National Institute of Health (grant identifier NIGMS-R01-GM063796 to G.A.V.), the Medical Research Council (grant identifiers MR/L00870X/1 and MR/L018578/1 to J.S.R.), and the European Union Seventh Framework Programme (grant identifier FP7-PEOPLE-2012-CIG: 333955 to J.S.R.). J.J.M. is grateful for support from the Carlsberg Foundation in the form of a postdoctoral fellowship (grant identifiers CF15-0552, CF16-0639, and CF17-0783). Several people are acknowledged for helpful discussions and providing analysis scripts including Jen Hsin, Ruibin Liang, and Alexander J. Pak. Computation was performed at the Research Computing Center at The University of Chicago (Midway and Makena machines), and through the Extreme Science and Engineering Discovery Environment (XSEDE) network: Stampede machine at the Texas Advanced Computing Center, The University of Texas-Austin and Comet machine at the San Diego Super Computing Center, University of California, San Diego. Furthermore, this research is part of the Blue Waters sustained-petascale computing project, which is supported by the National Science Foundation (awards OCI-0725070 and ACI-1238993) and the state of Illinois. Blue Waters is a joint effort of the University of Illinois at Urbana-Champaign and its National Center for Supercomputing Applications.

## AUTHOR CONTRIBUTIONS

All authors (J.J.M., J.M.A.G., J.S.R., and G.A.V.) contributed to research design, analysis of results and critical revision of the manuscript. In addition, J.J.M. and J.M.A.G. performed the theoretical and computational research, J.S.R. performed the experimental research, and J.J.M. wrote the first draft of the full manuscript.

